# Evidence for optimal behavior of predators from parallel field investigations in two distinct wolf-prey systems

**DOI:** 10.1101/2024.06.06.597612

**Authors:** Christina M. Prokopenko, Katrien A. Kingdon, Daniel L.J. Dupont, Taylor M. Naaykens, John Prokopenko, Julie W. Turner, Sana Zabihi-Seissan, Eric Vander Wal

## Abstract

Animals behave ‘optimally’ when they minimize their costs while maximizing their energetic gain. Optimal foraging theory predicts that with decreasing resource abundance, animals will increase 1) niche breadth, 2) territory size and movement distance, and 3) time spent at resource patches. We test these predictions by investigating clusters from GPS collared wolves (*Canis lupus*) in two predator populations with marked differences in their prey composition and abundance. As expected, wolves in a less abundant system increase niche breadth, territory size, step lengths, and time spent at each kill. Our work provides evidence of optimal behavior in an apex predator which can support population resilience across changing landscapes.

## Introduction

Animals, whether populations, groups, or individuals, exist across a gradient of resource abundance. Resource spatial distribution influences animal space use as animals evaluate the energetic gains from the resources against the costs to access the resources (Charnov 1976, Ostfeld 1990). Apex predators that hunt and kill large-bodied prey incur a high initial energetic cost during the hunting stage (Williams et al. 2014) and receive a high energy gain during consumption. A useful framework to evaluate predator decisions in diverse systems is optimal foraging theory (Prokopenko et al. 2023). Optimal foraging theory predicts that with lower resource abundance, animals will generalize their diet and expand their territory to meet energetic needs (Ford 1983). In addition, due to the increased movement costs between sparse prey, predators will spend more time at a kill (Charnov 1976). We apply these foundational expectations to behavioral observations of two predator populations with marked differences in their prey composition and abundance.

Dietary niche classifies animals by the proportion of food items consumed and is largely constrained by the resources available to an animal. Animals are either generalists, which consume many resource types equally, i.e. broad niche, or specialists, which focus on particular resource types, i.e. narrow niche (Pianka 1969). Whether a predator opts to specialize or generalize its diet can depend on the density of the preferred prey type. Within a given population, animals have been shown to alter their dietary niche across years or seasons to reflect fluctuations in primary prey availability (Burstahler et al. 2016) or when environmental conditions improve the success rate in hunting an alternative prey (Mysłajek et al. 2021). Species with broad food preferences, or those that consume alternate food types, will have wider dietary niches when preferred resources are scarce (Stephens and Krebs 1986). Generalist species have also been shown to have larger range sizes and therefore should occur in more patches across a landscape than species with narrow niches, i.e. specialists (Slatyer et al. 2013).

Animals construct their dietary niches within their home ranges, which are the space used to fulfill an organism’s needs to survive and reproduce (Burt 1943). The size of ranges is shaped by the amount of resources contained within, where increases in resources decreases range size (Ford 1983, Rizzuto et al. 2021). In a study across western Canada, wolves (*Canis lupus*) had smaller territories in more productive areas assumed to have more prey (Dickie et al. 2022).

Even at a fine scale, individuals adjust their range sizes as resources change temporally. Following a wildfire and subsequent resource pulse, black bears (*Ursus americanu*s) reduced their home range size (Crabb et al. 2022). In contrast to range contraction, expansion of home ranges increases costs of moving between resource patches and defending larger territories (Sells et al. 2021). Landscape variation in research distribution alters the search rate to meet the same energetic threshold (Mitchell and Powell 2007). In response to increases in resource availability, animals decrease their home range size to increase their exploitation efficiency.

Within home ranges, animals will adjust their activity budgets to be more efficient. The time spent in patches is influenced by the energy spent traveling between, searching within patches, and the energy consumed within a patch (Charnov 1976). For a predator, consuming a carcass (i.e. handling time) can be likened to patch residency. Accordingly, increased patch residency via handling time can offset travel costs between patches when prey are sparse. For example, wolves exhibited longer handling times as annual prey biomass per wolf decreased (Gable et al. 2023). Partial consumption of prey is an optimal foraging strategy when resource availability is high, the cost of searching is low, and predators are often satiated; wolves consumed smaller amounts of prey when kill rates were higher (Vucetich et al. 2012). At a fine scale, predators can decrease their handling of killed prey when more resources are available.

We studied the diet and behavior of two populations of a generalist predator, wolves, in different resource conditions. Overall, we expect wolves in the two populations to respond economically to their environment following optimal foraging theory resulting in predictable variation in behavior. We determine the influence of prey biomass (energy availability) on key population level patterns of niche breadth, time budgets, and range size of wolves via concurrent tracking and cluster investigation. We predicted that wolves with less prey available will respond by increasing 1) niche breadth, 2) home range size and movement distance, 3) time spent at patches through longer total times at kill clusters and higher revisitation to previous kills.

### Study Area

#### Riding Mountain National Park

Riding Mountain National Park (RMNP) is a 3,000 km^2^ federally protected area in southwestern Manitoba. RMNP is in Treaty 2 Territory, the ancestral lands of the Anishinabe people and the homeland of the Métis Nation (Figure 1A). Parks Canada works with First Nations from Treaties 2, 4, and 1 including Keeseekoowenin Ojibway First Nation, Ebb and Flow First Nation, Waywayseecappo First Nation, Rolling River First Nation, Tootinaowaziibeeng First Nation, Gambler First Nation, and Sandy Bay Ojibway First Nation. The border of the protected area is conspicuous against the surrounding agriculture-dominated landscape. The habitat within the park is a confluence of aspen parkland, mixed wood and boreal forest, and grassland prairie. There is a gradient in human activity decreasing generally northward, and westward. There is a network of trails, campsites, and two roads in the park. Most of the human activity is concentrated at the town site along the south shore of Clear Lake, which is outside of the core study area. Hunting is permitted immediately outside of the park boundary, and agricultural land use creates an abrupt land cover change.

**Figure 1.**
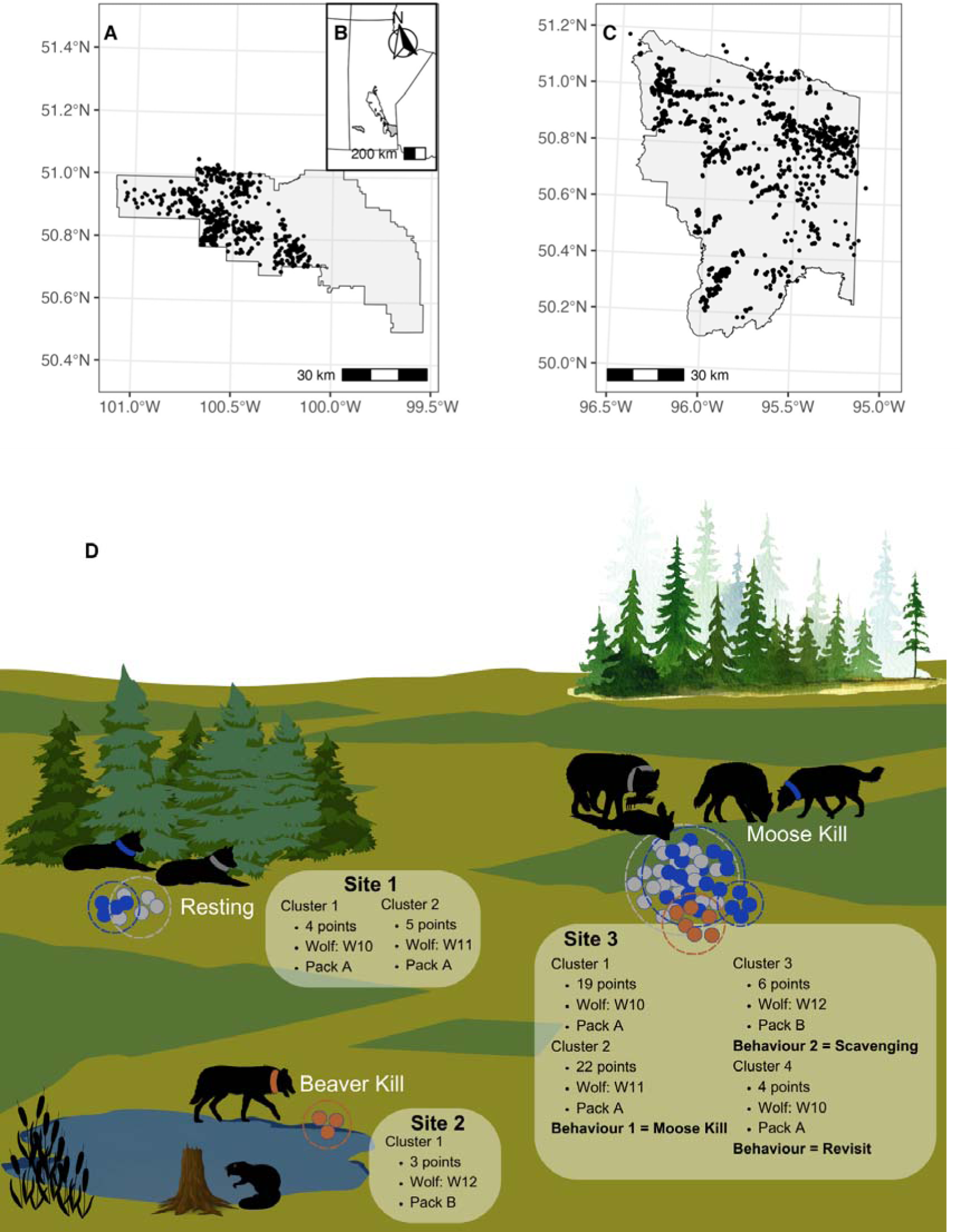
Cluster investigations were completed in two wolf populations A) RMNP with inset of B) location of both study areas within Manitoba and C) GHA 26. Borders of RMNP and GHA 26 filled in grey with the centroid locations of all investigated clusters in black; D) Conceptual diagram outlining the hierarchical structure of sites, clusters and behaviors investigated and determined through field work. Sites could be composed of multiple GPS clusters from a single or multiple wolves. Clusters were generated from the GPS locations (n>2) of a single wolf. Field investigations of clusters revealed primary and secondary behaviors associated with a site that could be linked to cluster characteristics.

Wolves are the apex predators in this system. During the present study the wolf population was approximately 71 individuals (consistently 60-80 individuals, 23 individuals/1000 km^2^). In addition to an abundant black bear population, there are populations of coyote (*Canis latrans*), lynx (*Lynx canadensis*), and incidental sightings of cougars (*Puma concolor*) (though no confirmed breeding population). The prey base for wolves is diverse and abundant in this area including moose (*Alces alces*), elk (*Cervus canadensis*), white-tailed deer (*Odocoileus virginianus*), beaver (*Castor canadensis*), and snowshoe hare (*Lepus americanus*). Historically, elk were more abundant than moose with populations fluctuating between 2000-6000 individuals. A significant population decline began in the late 1990’s for both species but was more substantial for elk, with moose becoming the most abundant ungulate in the park. During the study period, the populations of ungulates were estimated at around 2400 moose, 1200 elk, and 750 white-tailed deer. Recent beaver cache surveys conducted semi-annually show an increasing trend from an estimated 1700 beaver caches in 2013 to 2600 in 2016 (Parks Canada Report 2016).

#### Game Hunting Area 26

Game Hunting Area 26 (hereafter GHA 26) is a ∼7,200 km^2^ provincial management unit, located in southeastern Manitoba. GHA 26 is bordered by Lake Winnipeg to the west, Ontario to the east, Wanipigow River to the north, and Winnipeg River to the south (Figure 1C) and is part of the Boreal Shield Ecozone (Ecological Stratification Working Group 1995), which extends beyond the borders of the GHA designation and creates a continuous landscape. The landscape in GHA 26 is composed predominantly of coniferous and mixed forests, interspersed with rock outcrops, rivers, lakes, and bogs.

The study area is in Treaty 1, 3 and 5 Territory, the ancestral lands of the Anishinabe peoples and the homeland of the Métis Nation. The study area includes the traditional territories of Sagkeeng, Black River, and Hollow Water First Nations, Brokenhead Ojibway Nation, and Northern Affairs communities (Bissett, Aghaming, Manigotagan, and Seymourville). The area is predominantly public land, and GHA 26 is used by many different user groups for both recreational purposes and resource development. There are two provincial parks in the area: Nopiming Provincial Park and Manigotagan River Provincial Park. The local economy is predominantly based on resource extraction, including past forestry activity, which has led to a network of roads including primary roads, logging roads, trails, and hydroelectric transmission line rights-of-way (ROW).

Wolves are the apex predator in this system with an average population of 128 individuals between 2012 and 2018 (range 106-141 individuals, ∼17 wolves/1000km^2^). The predator guild includes populations of black bears, coyotes, and lynx. The prey base for wolves in GHA 26 is diverse, yet spatially distinct, with moose found primarily in the northern half of the study area and white-tailed deer found primarily in the southern half of the study area. Secondary prey species include beavers, snowshoe hare, and woodland caribou (*Rangifer tarandus caribou*). Moose were historically found throughout the study area, but are now found primarily in the northern half, which has led to a reduction in moose abundance and distribution in the study area. The primary prey species for wolves in GHA 26 are moose, with an estimated population of 2400 individuals in the early 2000s but experienced a drastic decline by approximately 50% in the early 2010s. The most recent moose survey conducted in 2020 provided a population estimate (unadjusted for sightability) of approximately 1200 moose. A beaver cache survey was conducted in 2018 and the cache density throughout the study area was estimated at 1,14 beaver caches per km2.

## Methods

Detailed methods of data cleaning, niche breadth calculations, movement and home range analysis are in section S1 of the supplemental file. Data and code (Python version 2.7.1, R version 4.3.0) and are available on GitHub https://github.com/CMProkopenko/wolf_clusters and will be archived on Zenodo.

### Cluster Investigations

Wolves were collared with GPS telemetry collars in RMNP (25 individuals) and GHA 26 (38 individuals) between 2014-2018 following Memorial University AUP 16-02-EV (Packs collared by year in supplemental file S2). Continuous and extensive fieldwork investigations determined the timing and location of wolf behaviors, including wolves killing prey, scavenging, denning, and resting (see Table 1 for full description of behaviors). Increased density of GPS locations, i.e., ‘clusters’, indicated important areas of wolf activity (Figure 1C). To identify clusters, we used an algorithm in Python, which was presented in Knopff et al. (2009), created by Warren (2008), and adapted for wolves (Webb et al. 2008, DeCesare 2012, Irvine et al. 2022). In our study, we defined a cluster as GPS locations that occurred within a 300m radius and 96 hours of a previous location. This cluster algorithm identified 10 628 clusters from all collared wolves in GHA 26 between February 2014 and March 2020 and 6323 clusters in RMNP (Feb 2016 to Jan 2018).

**Table 1.**
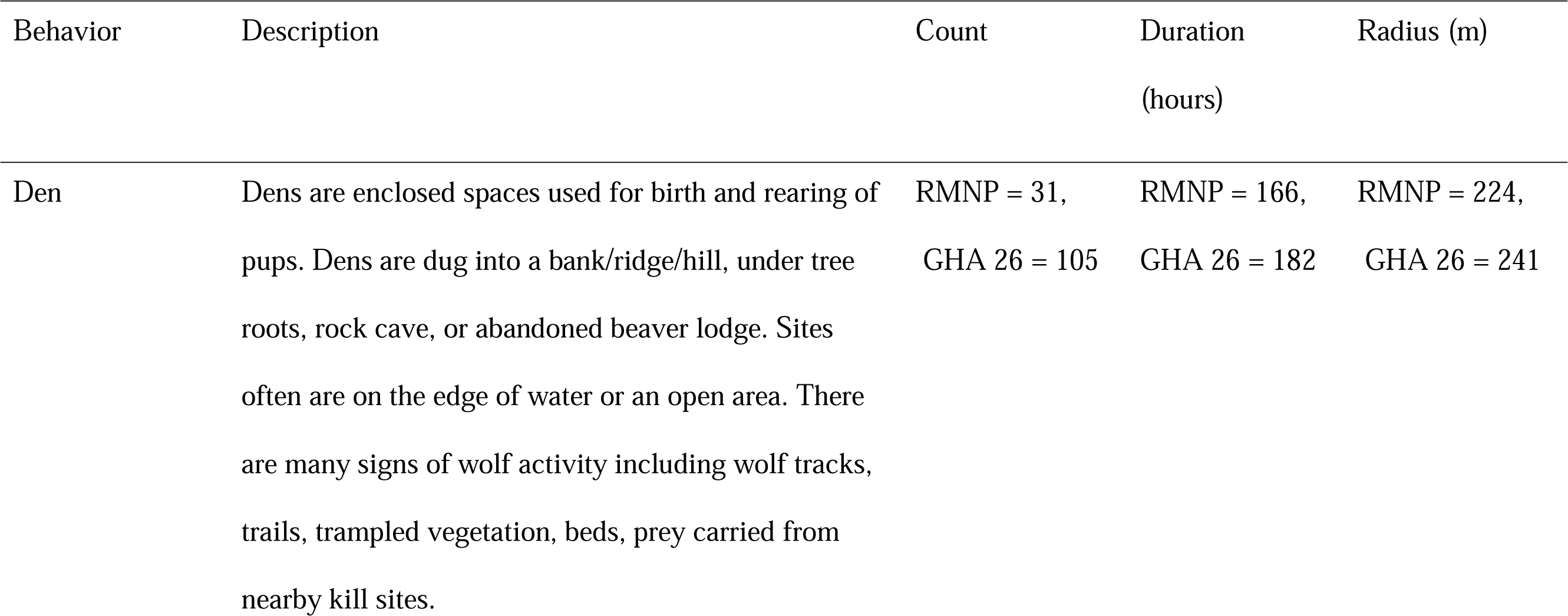
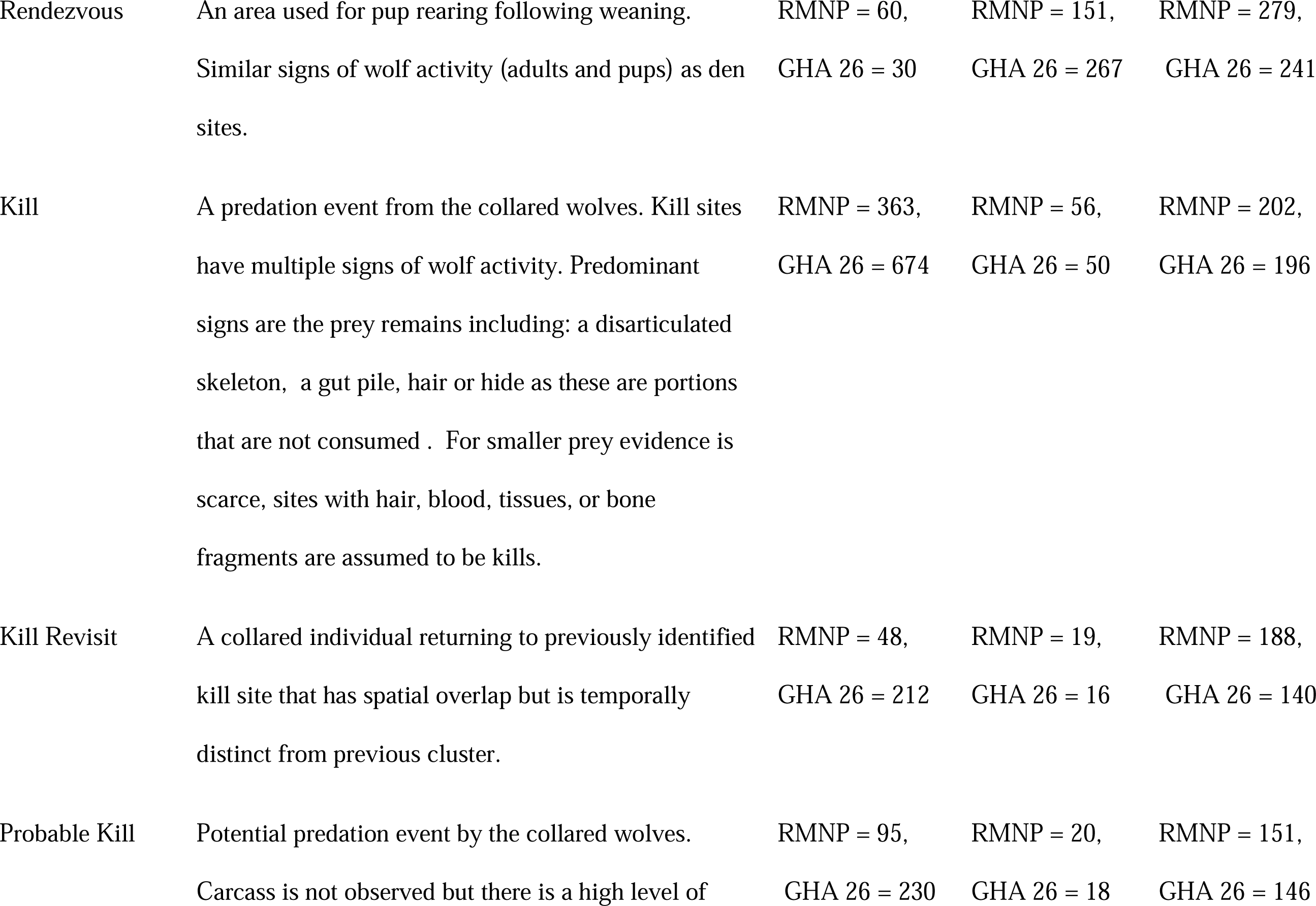
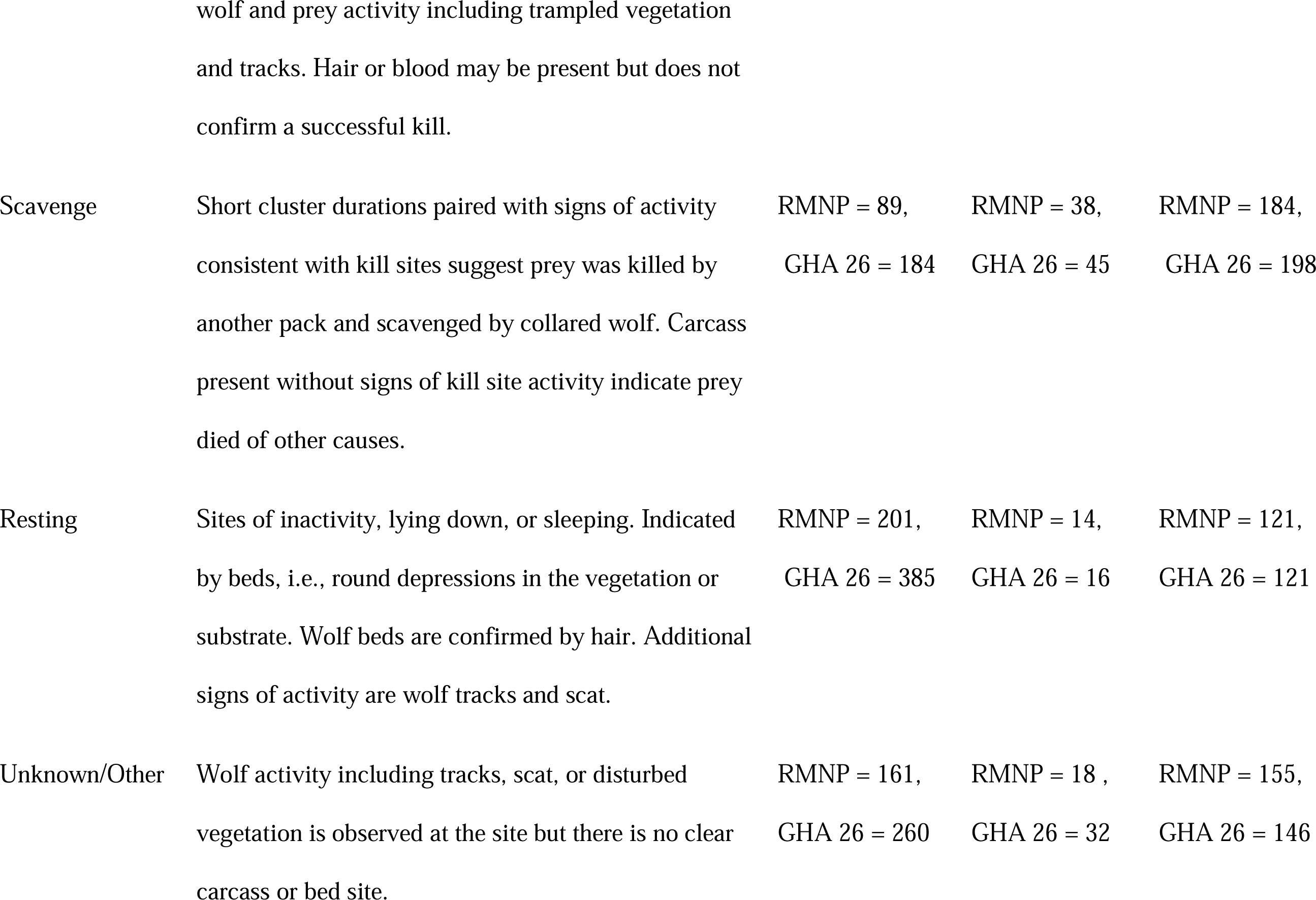
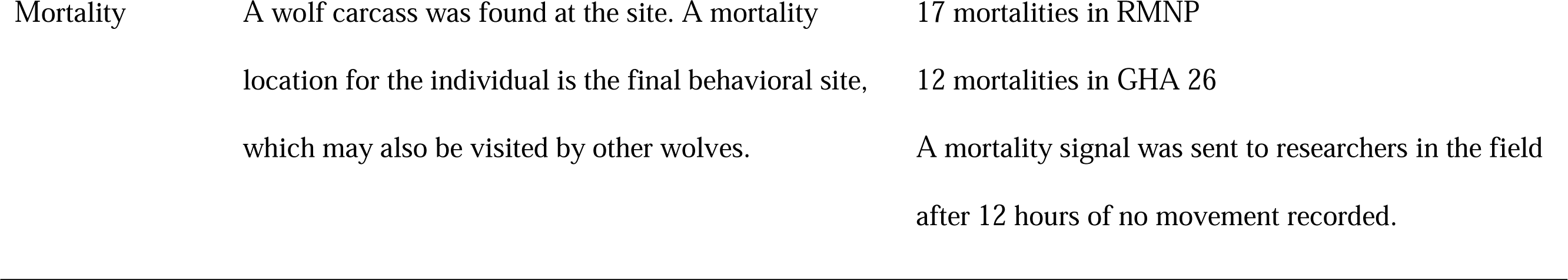
Description of behaviors assigned to clusters during field investigations. Behaviors are presented in order of the hierarchy that was determined during the cleaning phase of our process. It is common for behaviors lower in the hierarchy to occur at sites with higher ranked behaviors. For example, most kills often include resting and scat activity. The total count of clusters, the median duration in hours, and median radius in meters of these clusters are presented by study area.

To direct our sampling effort, we categorized clusters into three classes created based on the total number of locations in the clusters (small = 2-4 locations, medium = 5-7 locations, large = 8+ locations). Clusters were most often accessed via snowmobile, quad, or occasionally helicopter, as well as by canoe or boat in GHA 26 and horseback in RMNP. We searched for physical evidence of wolf behavior starting at the geometric center of the cluster moving outwards from this point. We investigated a 15 m radius around each GPS fix in the cluster to account for GPS error. On average, we investigated clusters within a mean of 13.8 days of the first location occurrence (median = 9 days).

We identified unique behaviors within a single cluster corresponding to the physical evidence present at clusters (Table 1). We then termed these clusters as various ‘sites’ corresponding to the primary behavior identified (Figure 1C). For example, we classified clusters as kill sites when a carcass was present coupled with a high degree of activity indicative of an interaction between predator and prey. In some instances, we determined clusters with carcasses were instead the result of scavenging based on evidence that the animal died due to another cause, such as lack of wolf activity, or if the collared wolf spent too short of a duration at the site to have been responsible for the death of the animal, e.g., < 2 hours at a large ungulate kill. We determined prey species and for ungulate prey we determined sex and age (adult vs. calf) based on size, dentition, and hair pattern. At the site we collected biological samples (prey hair, bone marrow, wolf scat). In the absence of a carcass, we used other physical evidence from wolves or other species to determine the behavior that occurred at the site such as tracks, hair, beds, scat, and dens. For example, beds under a spruce tree containing wolf hair were indicative of a resting site. A single site could have more than one behavior, for example if we observed both kill and resting evidence. However, we would consider “kill” to be the primary behavior and “resting” to be the secondary. For the data cleaning protocol following fieldwork, see supplemental file S1.

### Scat Dissection

We collected scat samples during cluster investigations at sites and opportunistically when traveling to sites in the field. We typically collected only the first sample we came across unless there were multiple prey found at the site or the samples were visually distinct. In GHA 26, we collected 531 scat samples from February 2014 to March 2020, while in RMNP we collected 296 scat samples from Feb 2016 – Jan 2018. Scats were stored for a minimum of 2 weeks at -80°C before handling to kill any potential parasites. Scats were manually strained through a sieve to separate hair and other undigested remains from fecal matter. Twenty hairs were arbitrarily selected from each scat for identification. Hair color, size, and cuticular scale pattern were used to identify prey remains(Adorjan and Kolenosky 1969, Kennedy and Carbyn 1981). This method allowed for distinctions to be made between adult and juvenile ungulate hairs. We evaluated potential bias for large clusters by comparing the number of locations across all clusters investigated. We summarized the number of locations, duration, and radius for clusters by behavior and kill clusters by prey type.

### Diet Analysis

We calculated the composition of wolf diet as both percentage of occurrence in scat and at clusters. This was calculated as either the percentage of scats containing a prey species relative to the total number of scats or as the percentage of clusters where a prey species was either killed, scavenged upon, or was likely to have been consumed (i.e., probable kill, Table 1) relative to the total of these cluster behaviors.

We compiled all scat and cluster data to compare wolf diet composition between RMNP (2016-2017) and GHA 26 (2014–2019). As not all prey species are equally available across seasons, we calculated niche breadth (*B,* Levins 1968) for both snow (November to April) and snow free (May to October) seasons. We classified wolf prey into the 4 general groups. The B index, therefore, could range from 1, indicating a strong specialization to one group of prey, to four, indicating equal preying on all groups.

### Space-Use Analysis

We quantified home ranges for each individual wolf in both snow and snow free seasons using 95% autocorrelated kernel density estimators (aKDEs; Fleming et al. 2015, Noonan et al. 2019) using the amt (Signer et al. 2019) and ctmm (Calabrese et al. 2016) packages in R.

## Results

### Cluster Investigations

In both RMNP and GHA 26, we investigated 16% of the total GPS clusters (RMNP: 1061/6433 clusters; GHA 26: 1897/11804) generated during the study periods. We identified both the primary and secondary behaviors at each cluster and calculated various spatial and temporal metrics that could provide a link between behavior and remotely sensed data.

Wolf GPS clusters ranged in size from 2 to 275 locations. Small clusters (2-4 locations) occurred more frequently, and we visited a higher total number but smaller proportion of small clusters (Figure 2A, 2B, and 2C). Investigation rates increased with the number of locations and peaked around a proportion of 0.7 for clusters that had 20-60 locations, then declined as cluster size continued to increase (Figure 2C). Reproductive behaviors, den and rendezvous were associated with the largest clusters (Table 1, Figure 2D and 2E). In comparison, resting clusters had some of the fewest locations and duration (Table 1, Figure 2D and 2E).

**Figure 2.**
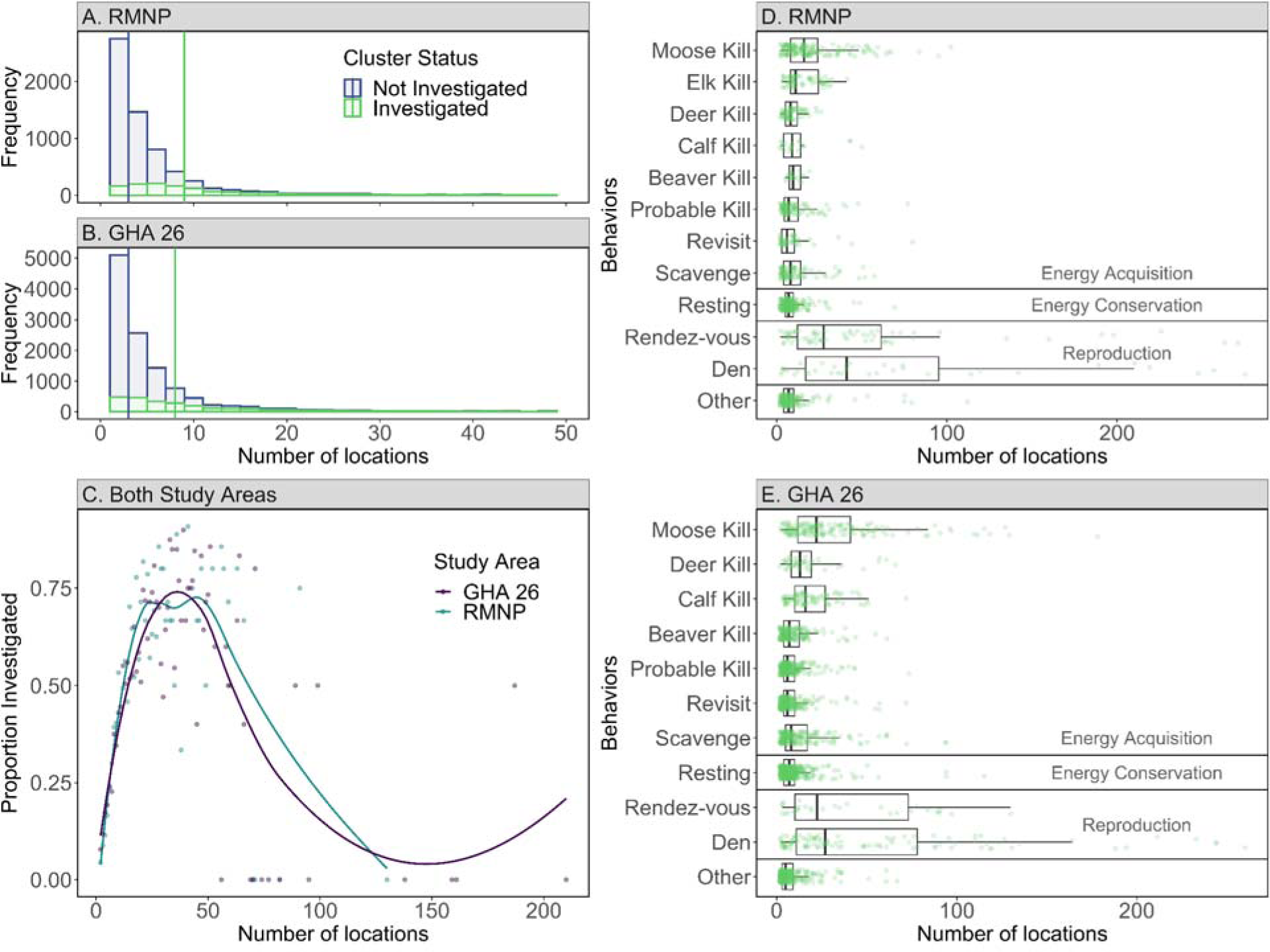
Cluster size, defined by the number of GPS locations (2-hour schedule) included in the clusters investigated and not, proportion investigated, and behaviors at the clusters for Riding Mountain National Park investigations from 2016 to 2017 and for Game Hunting Area 26 from 2014 to 2019. Histograms depicting the count of clusters that were investigated and not investigated by the number of locations for A. RMNP and B. GHA 26. Vertical lines indicate median number of locations for all clusters investigated (9 locations for RMNP, 8 locations for GHA 26) and not investigated (3 locations for both study areas) Note: x-axis was limited to 50 locations to remove long tail of 0 or 1 occurrence extending to 200 locations. C. Scatterplot and line (loess smoothing method in ggplot2) showing the relationships between proportion of clusters investigated and the number of locations in clusters for both study areas. Boxplots and points of behavior categories at clusters of different sizes for D. RMNP and E. GHA 26. Primary behaviors and prey were used to define the categories on the y axes. Horizontal lines divide behaviors further from top to bottom into energy acquisition, energy conservation, reproduction, or other. Definitions of behaviors with key signs observed at clusters during investigations are in Table 1.

The dominant primary behavior identified at clusters during investigations was kill (363 clusters in RMNP, 674 in GHA 26), followed by resting (201 clusters in RMNP, 385 in GHA 26) for both study areas (Table 1). Time spent at kill clusters varied between prey species and study areas (Figure 2, Table S2.1). In RMNP, wolves spent the most time on average at large ungulate kills including moose (68 hours) and elk (60 hours) but less time at small ungulate kills such as white-tailed deer (22 hours) and calves (26 hours). In contrast, wolves in GHA 26 spent longer at kills of most ungulate prey including moose (92 hours), calves (63 hours), and white-tailed deer (22 hours). However, wolves spent less time at beaver kills in GHA 26 with clusters lasting a median of 22 hours compared to 46 hours in RMNP. GHA 26 wolves had more kill revisits than RMNP (Table S2.1).

Wolves in the same pack often simultaneously visit the same site or departure/return events by a single wolf can lead to generating multiple clusters that overlap in space and time. For example, a single moose kill can be associated with 4 individual clusters if multiple wolves visit the site or if a wolf leaves and returns (Figure 1D). To further summarize wolf behavior in these two regions, we categorized unique behaviors and feeding events as sites, using the primary and secondary behaviors observed across all clusters associated with these unique events. We identified 549 unique sites from the 1061 clusters we investigated in RMNP and 1091 sites from the 1897 investigated clusters in GHA 26.

### Diet composition and niche breadth

We summarized wolf feeding behaviors at GPS clusters into unique kill, probable kill, and scavenge sites. Wolves in both RMNP and GHA 26 fed on a wide variety of species (Table 3). Both scat analysis and site investigation revealed over 90% of wolf diet consisted of adult ungulates (RMNP: elk, moose, white-tailed deer; GHA 26: moose, white-tailed deer), ungulate calves, and beavers, as well as snowshoe hares in GHA 26. Niche breadth calculations indicated that wolves from RMNP had a more specialized diet compared to wolves in GHA 26 (Table 2). Overall, in GHA 26, wolves consumed fewer adult ungulates, but more beavers, hares, and ungulate calves to increase the dietary niche breadth.

**Table 2.** Diet composition and niche breadth calculations of wolves in Riding Mountain National Park (2016-2017) and Game Hunting Area 26 (2014-2019) from scat samples and at investigated sites identified as kill, probable kill or scavenge sites. Composition is reported as the percent occurrence, as well as total samples in brackets, calculated for each data source and study site. Dashed lines (-) indicates the species is not found in that region. We analyzed dietary niche breadth calculations year-round and separately for snow (November-April) and snow-free (May-October) seasons.

|  | RMNP |  | GHA 26 |  |
| --- | --- | --- | --- | --- |
|  | Site | Scat | Site | Scat |
| Season | Dietary Niche Breadth |  |  |  |
| Snow | 1.18 | 1.43 | 2.95 | 2.14 |
| Snow-Free | 2.54 | 2.52 | 2.20 | 3.28 |
| Year-Round | 1.54 | 1.80 | 2.66 | 2.82 |
| Species | Diet Composition |  |  |  |
| Moose Adult | 44.91 % (119) | 34.39 % (54) | 21.27 % (151) | 33.39 % (185) |
| Deer Adult | 16.98 % (45) | 10.19 % (16) | 9.58 % (68) | 11.91 % (66) |
| Elk Adult | 15.47 % (41) | 27.39 % (43) | - | - |
| Unknown Ungulate Adult | 2.26 % (6) | 0 | 0 | 0 |
| **Adult Ungulate Total** | **79.6 % (211)** | **71.97 % (113)** | **30.8 % (219)** | **45.3 % (251)** |
| Moose Calf | 2.64 % (7) | 1.91 % (3) | 10.28 % (73) | 7.4 % (41) |
| Deer Calf | 0.75 % (2) | 0 | 0.42 % (3) | 0.54 % (3) |
| Elk Calf | 0 | 0.64 % (1) | - | - |
| Unknown Ungulate Calf | 2.26 % (6) | 0 | 0 | 0 |
| **Calf Ungulate Total** | **5.66 % (15)** | **2.55 % (4)** | **10.7 % (76)** | **7.94 % (44)** |
| Beaver | 10.19 % (27) | 17.83 % (28) | 42.11 % (299) | 32.67 % (181) |
| Snowshoe Hare | 0 | 0 | 9.3 % (66) | 3.61 % (20) |
| **Medium Mammals** |  |  |  |  |
| **Total** | **10.19 % (27)** | **17.83 % (28)** | **50.6% (381)** | **37.48% (187)** |
| Grouse | 0.75 % (2) | 0 | 1.55 % (11) | 0 |
| Goose | 0.38 % (1) | 0 | 0.28 % (2) | 0 |
| Duck | 0 | 0 | 0.14 % (1) | 0 |
| Raven | 0 | 0 | 0.14 % (1) | 0 |
| **Bird Total** | 1.43% (4) | 0% (0) | 2.12% (16) | 1.62 % (9) |
| Marten | 0 | 0 | 0.14 % (1) | 0 |
| Mink | 0.38 % (1) | 0 | 0 | 0 |
| Muskrat | 0 | 0 | 0.28 % (2) | 0 |
| Unknown Rodent | 0 | 3.18 % (5) | 0 | 2.17 % (12) |
| **Small Mammal Total** | 0.38 % (1) | 3.18 % (5) | 0.40% (3) | 2.17 % (12) |
| Bear | 0.75 % (2) | 1.27 % (2) | 1.97 % (14) | 1.62 % (9) |
| Fish | 0 | 0 | 1.69 % (12) | 3.43 % (19) |
| Vegetation | 0 | 3.18 % (5) | 0 | 0 |
| Human Disposal | 1.14 % (3) | 0 | 0.53% (4) | 0 |
| Unknown | 1.13 % (3) | 0 | 0.28 % (2) | 1.62 % (9) |

Comparing the biomass for the top five prey species in each study area shows moose constituted the highest proportion of wolf diet across study area, sample type and season (Figure 3). In RMNP, we observed similar diet proportions in scat samples for moose and elk in snow free seasons, with increased disparity during snow seasons (Figure 3). Cluster data revealed beaver and calves made up a higher percentage of the diet in snow free seasons. In GHA 26, scat biomass data indicated a seasonal shift in percentage of moose in wolf diet that was not as strongly observed in the cluster data. The percentage of beaver biomass increased in snow free months, particularly in scat samples. We observed a seasonal shift in diet across almost all packs in GHA 26 (Figure 4) where beaver and calves comprised a much higher percentage of wolf diet in snow free months. There were fewer observable seasonal shifts in wolf diet in RMNP. Beaver diet percentage was much lower compared to GHA 26 overall, but we observed an increase in calf percentage in snow free seasons.

**Figure 3.**
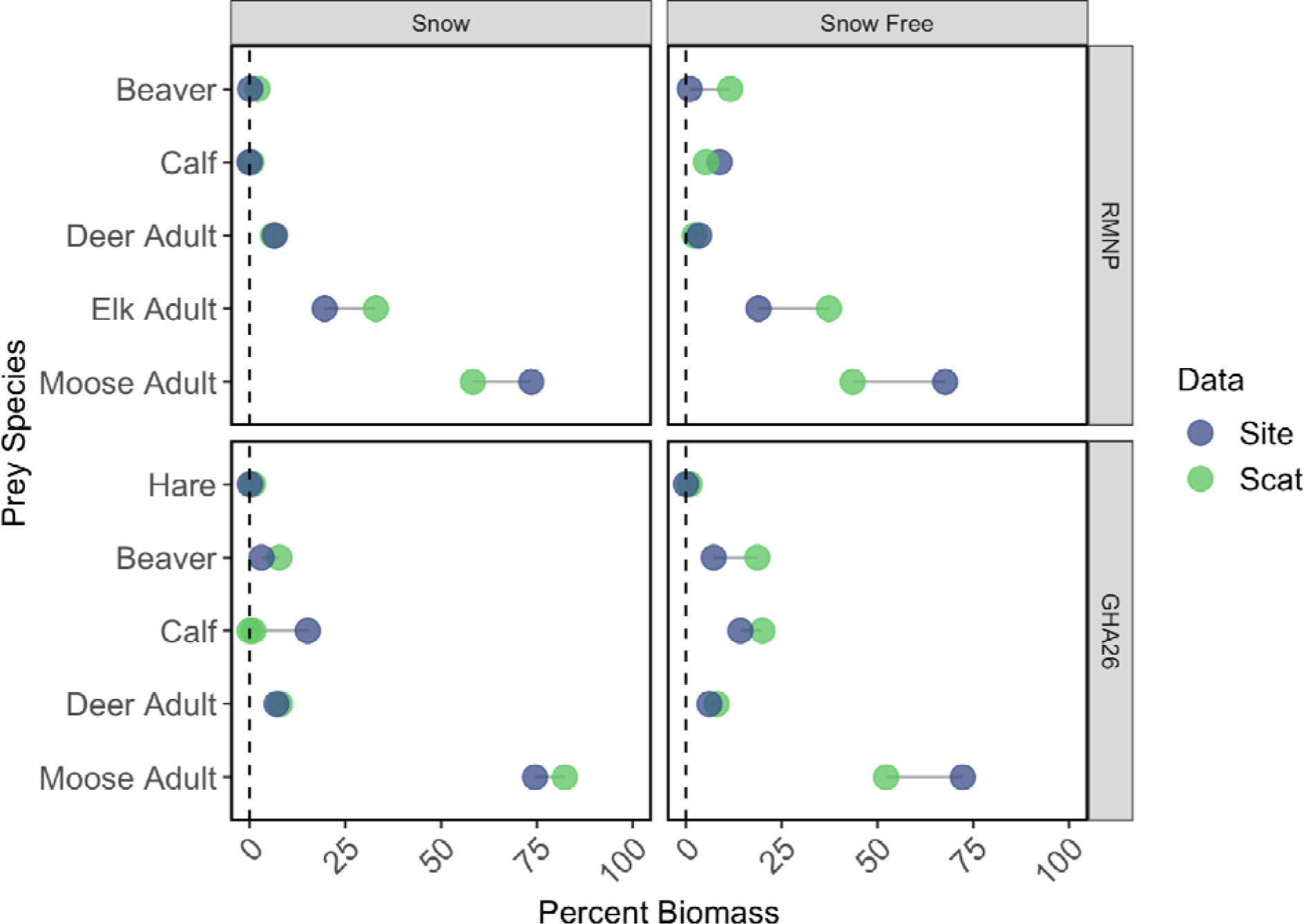
Seasonal percent biomass contribution of prey type to gray wolf diet in RMNP (2016 - 2017) and GHA 26 (2014-2019) determined through two data sources; kill site investigations of clusters from collared wolves (i.e., Site) or found in wolf scat samples collected at all site types (i.e., Scat).

**Figure 4.**
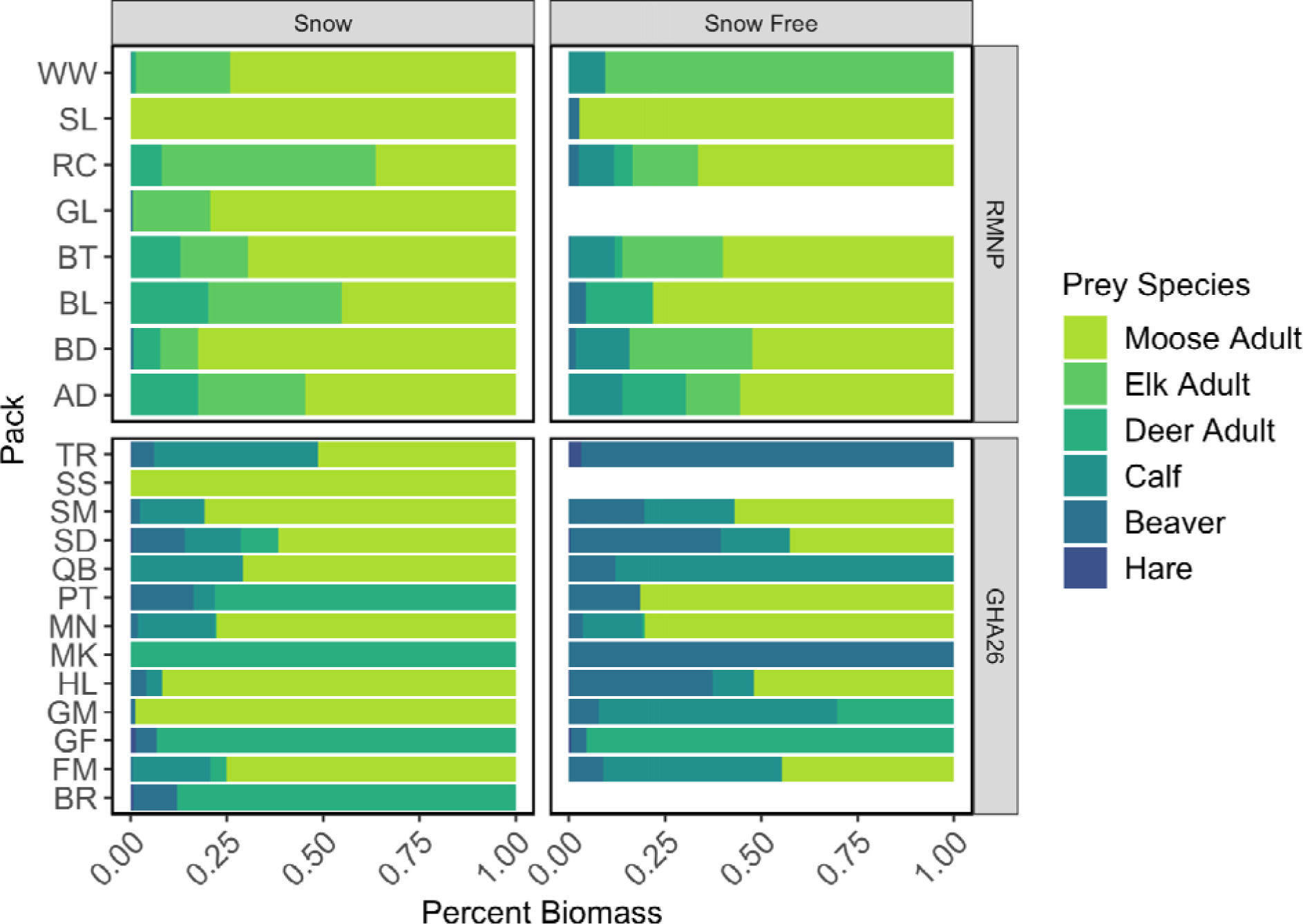
Seasonal percent biomass contribution of prey type to gray wolf diet across packs in RMNP (2016 - 2017; n = 8) and GHA 26 (2014-2019; n = 13) determined through kill site investigations of clusters from collared wolves.

**Figure 5.**
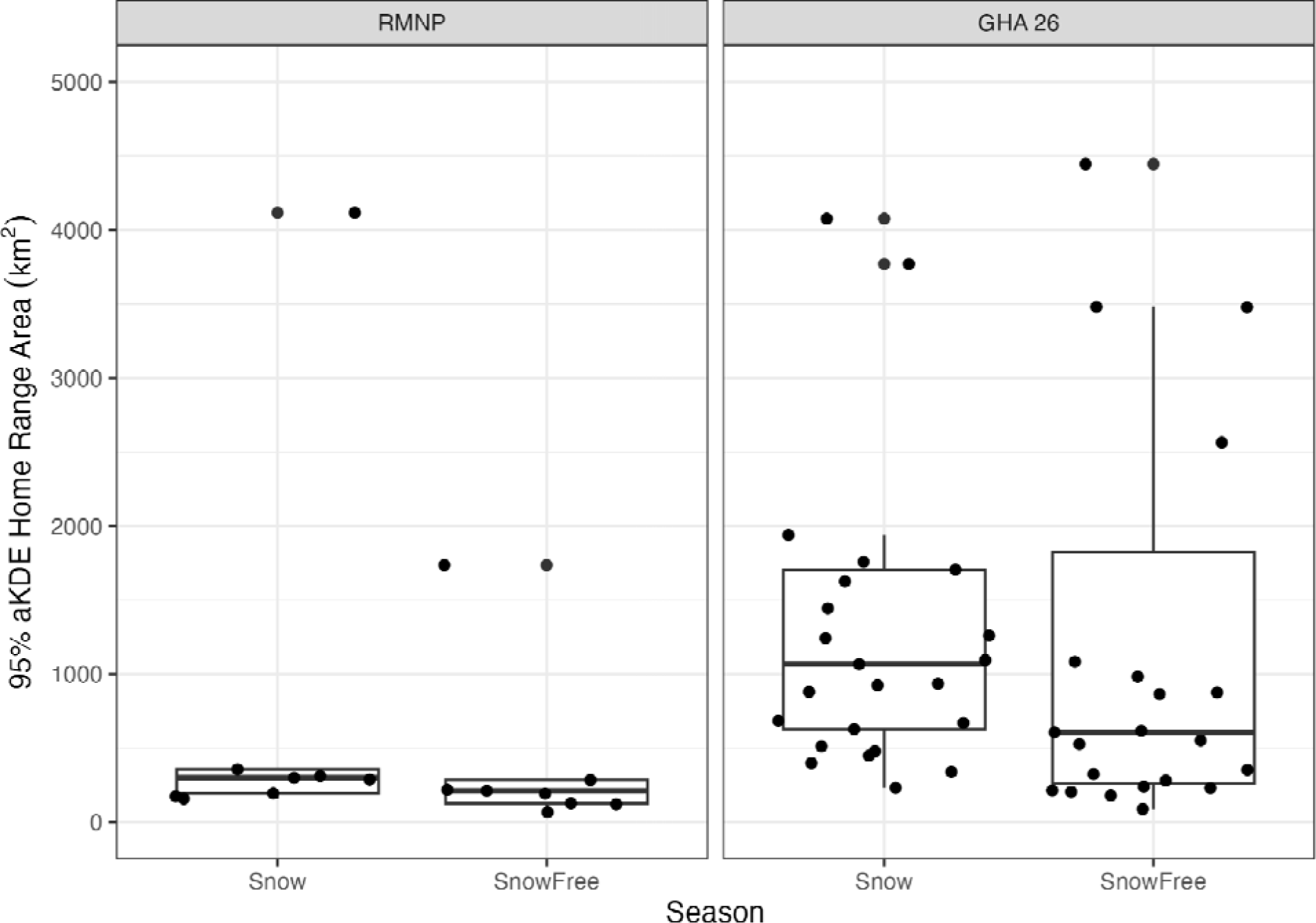
Home range areas (km2) of gray wolves in RMNP (2016 - 2017; n = 8) and GHA 26 (2014-2019; n = 13) calculated from 95% aKDE isopleths. Home range sizes greater than 5000 km2 were included in the boxplot calculations but not included in the figure for visualization purposes. See supplemental file Table S2.2. for all values.

### Home range and step length

Across both study sites, wolf packs typically had smaller home range sizes during snow free seasons and wolves in RMNP had smaller home ranges than wolves in GHA 26 (Figure 4). In RMNP home ranges had a median area of 202.9 km^2^ (range: 67.4 to 1736 km^2^) in snow free seasons compared to 294.8 km^2^ (range: 159.1 to 4116.7 km^2^) in the snow seasons. Wolves in GHA 26 had median home range sizes of 552.0 km^2^ (range: 88.6 to 4445.9 km^2^) and 934.4 km^2^ (range: 233.0 to 4076 km^2^) in snow free and snow seasons respectively. RMNP wolves moved less over the 2-hour time periods (median = 125 m, mean = 793 m) than GHA 26 wolves (median = 154 m, mean = 1077 m), see supplementary file Figure S2.5. RMNP did not adjust their movement seasonally, compared to GHA 26 (Figure S2.6)

## DISCUSSION

In our comparative study of two populations, wolves increased their diet breadth, home range size and movement distances, and patch residency time in areas with less prey. Wolves in areas with lower resource availability were more generalized than wolves in the research rich areas. The shift to pulses in smaller bodied prey was greater in the low resource population compared to the high resource population. Time budgets between the two populations differed: wolves spent more time at kills in low resource areas. Wolves spatially responded by increasing movement rates and territory sizes with decreasing resource availability. We find responses at the population and pack level follow the fundamental expectations of animals attempting to maximize energy acquisition and minimize time expenditure supporting our predictions from optimal foraging theory.

Wolf niche breadth increased with decreasing prey abundance, as expected from optimal foraging theory. At a population-level, wolves in RMNP overlap with a higher density of ungulate prey populations than GHA 26. Wolves in RMNP consumed primarily large ungulate prey. In comparison, wolves in GHA 26 had a broader niche breadth and more generalized diet, consuming more calves and small mammals. Wolves have the potential to expand their diet when ungulate prey are not available to include diverse alternative prey, such as fish (Darimont et al. 2003) or marine prey (Roffler et al. 2021). In GHA 26, beavers were the most frequently consumed prey item, contributing 25% of consumed biomass in the snow free period, in contrast to RMNP that focused on large ungulates throughout the year. In many wolf systems, beavers are an important subsidy for wolves and can increase recruitment (Gable et al. 2018). Here, wolves adjusted their diet to smaller prey in the area with lower resources more than those in the high resource area, as expected from optimal foraging theory.

Optimal foraging predicts increased competition in areas of low resource abundance will drive individuals to add new items to their diet, widening their trophic niche, and forming a population of generalist foragers (Stephens and Krebs 1986). Alternatively, we could view this through competition theory, where increased competition leads to stable coexistence through niche differentiation, reducing dietary overlap between competitors (Schoener 1974, Pianka 1981). Both RMNP and GHA 26 both have black bear populations that are a potential competitor for wolves though in some instances were a prey item (Table 2). Competition and resource availability are not mutually exclusive but are inherently linked as a consumer selects resources. For example, caribou consume vascular plants when available but predation or competition push caribou towards consuming lichen (Webber et al. 2022). Assessing the various factors that contribute to the niche breadth within a species will strengthen our descriptions of critical habitat and help to predict distributions of animals.

Territories were larger in the study area with less abundant prey, supporting the idea that wolves expand their distribution to meet energetic needs. Similarly, territories of wolves across the boreal forest decreased with increasing resource density (Dickie et al. 2022); wolves in low resource conditions had territory sizes of less than half that of wolf territories in high resource conditions. In our study, year-round territory sizes of resident wolf packs spanned from 137.9 to 2621.7 km^2^ in RMNP. The territory sizes in RMNP fall below or at the low end territory size of precious estimates from wolves across the boreal (Dickie et al. 2022) indicating the resources in RMNP are higher. In GHA 26, territories were 213.9 to 12901.3 km^2^ indicating lower resource conditions than RMNP. In addition, step lengths (2hr scale) were shorter for RMNP than for GHA 26. Smaller territories come with reduced defense and movement costs (Sells et al. 2021), which we find with reduced movement distances in RMNP. Thus, the wolves in RMNP experience multiple energetic benefits compared to wolves in GHA 26.

In addition to resource availability, anthropogenic disturbance can alter space use behavior, but high resource density and low disturbance were co-occurring factors in our study; RMNP was an area with reduced disturbance and smaller territories compared to GHA 26. Increasing linear features on the landscape decreased wolf territories almost twice as much as increasing resources due to improved resource exploitation via movement efficiency (Dickie et al. 2022). Alternatively, linear features can fragment habitats and be perceived as a risk, which would in turn erode habitat quality. In support of this, wolves in Italy increased their territory sizes with road density (Mancinelli et al. 2018). Wolf space use behavior, specifically their territory size, was characteristic of differences resulting from resource abundance, but the high resource area coincided with being a protected area with a distinct border.

The time budgets of wolves differed between the two populations. Overall, RMNP wolves spent less time at large kills and scavenging than GHA 26 wolves. In addition, there were limited revisits of kills in RMNP, which we interpret as wolves opting for new kills instead of spending time at previous kills. This behavior aligns with the expectation from Marginal Value Theorem that consumers reduce their time within resource patches as resource abundance increases (Charnov 1976). A prey item could be compared to a ‘patch’ for carnivores. For example, wolves meeting their energetic needs reduced the amount of each prey item consumed and not the number of prey killed (Zimmermann et al. 2015). Indeed, partial prey consumption is an optimal foraging strategy that could be generating shorter times at kills in RMNP.

Handling times generally increase with the size of prey, yet there were important exceptions between species and study areas that do not follow this pattern. For example, RMNP tended to spend longer at beaver kills than white-tailed deer. The time at a kill includes both the time it takes to chase and subdue prey with consumption and digesting time. Wolves have been found to use alternative techniques when hunting beavers including sit and wait tactics associated with ambush hunting (Gable et al. 2021) However, given that some attempts lasted up to 30h (mean 4h; Gable et al. 2021), this is a costly tactic in a temporal budget. In support of this, RMNP wolves spent twice as long at beaver kills than GHA 26 wolves, a prey GHA 26 wolves consumed over twice as much as RMNP wolves. Beaver kills were the only incidence where RMNP wolves had a longer average time at kill sites. Future work in our system will focus on decomposing the predation sequence and calculating population metrics such as kill rates and predation rates.

We found evidence for all predictions at different scales of behavior, but there is likely a hierarchical organization to how animals respond to resource limitation which we did not test. For example, the behavioral response to low resources could begin with adjustments to movement rate and activity which propagates to changes in territory size. Future work could describe the sequence of animals adjusting their behavior as resources change. Further, we identified that the low resource population used the full suite of predicted responses, indicating that these animals may be approaching a threshold where no further behavioral adjustment is possible. Identifying the threshold of animal behavioral limitation brings these responses to their population-level consequences. Consumers can mediate changes in resources by adjusting their space use and time budgets to behave optimally by maximizing their energy gain and minimizing costs. However, wildlife are restricted by human disturbance (Tucker et al. 2018) and conflict increases when humans and wildlife overlap spatially (Teichman et al. 2013). Determining where and how consumers, specifically large carnivores, will survive with future changes can inform the management of these populations.

## Supporting information

S1 Methods Details S2 Additional Results

## Author contribution statements

Conceptualization for the paper was led by KK and CMP with support from DD and EVW. Field methods development was led by DD supported by SZS CMP and KK. Cluster investigations were led equally by CMP and SZS in RMNP and DD and KK in GHA 26. Data collection and processing was completed by all coauthors to various extents. JP and CMP lead the development of code to clean collar locations and clusters, and modified the cluster algorithm, Analysis and visualization was led by CMP and KK supported by JWT and DD. EVW supported coauthors through supervision and lead funding acquisition. CMP and KK lead the writing of the original draft and received comments from all coauthors.

## Acknowledgements

We respectfully acknowledge that RMNP is in the ancestral lands of the Anishinabe people and the Homeland of the Métis Nation, within Treaty 2 territory and Parks Canada works with First Nations from Treaties 2, 4, and 1. We would like to acknowledge that our research in Eastern Manitoba occurs within Treaty 3 and 5 Territory and within the Homeland of the Métis Nation. CMP was supported by a Vanier Canada Graduate Scholarship, KAK received an NSERC CGS-M and CGS-D Scholarship, TN received funding from Alice Chambers-Hyacinth Colomb Student Assistantship, EVW received a NSERC Collaborative Research and Development Grant. Funding and logistic support for this project was provided primarily by Parks Canada Agency (Riding Mountain National Park of Canada), with additional support from the Fish and Wildlife Enhancement Fund, and the Nature Conservancy of Canada. We thank T Sallows, KJ Kingdon, D Bergeson, R Robinson, R Baird, and R Grzela from Parks Canada and A Critchley and R Sandstrom from Shoal Lake Aviation for their contributions during data collection in RMNP. We are grateful for all the field technicians who assisted with data collection including S Graham and S Walmsley.The work in eastern Manitoba would not be possible without the support of Manitoba Wildlife and Fisheries Branch, Manitoba Hydro, the Fish and Wildlife Enhancement Fund, Lac du Bonnet Wildlife Association and local communities. In particular, we thank K Leavesley, V Harriman, K Rebizant, D Brannen, D Bulloch, and J Matthewson for their continued support. Thank you to I Lavoie, A Scott, S Graham, J Budzinski, S Böttcher, M Erich, C Clayton, M Ewacha, N Kaminski, R Bartel, and J Dupont for field support and many others. Members of the Wildlife Evolutionary Ecology Lab at Memorial University of Newfoundland and Labrador and the Wildlife Restoration Lab at University of British Columbia Okanagan graciously provided comments on earlier versions of this manuscript.

---

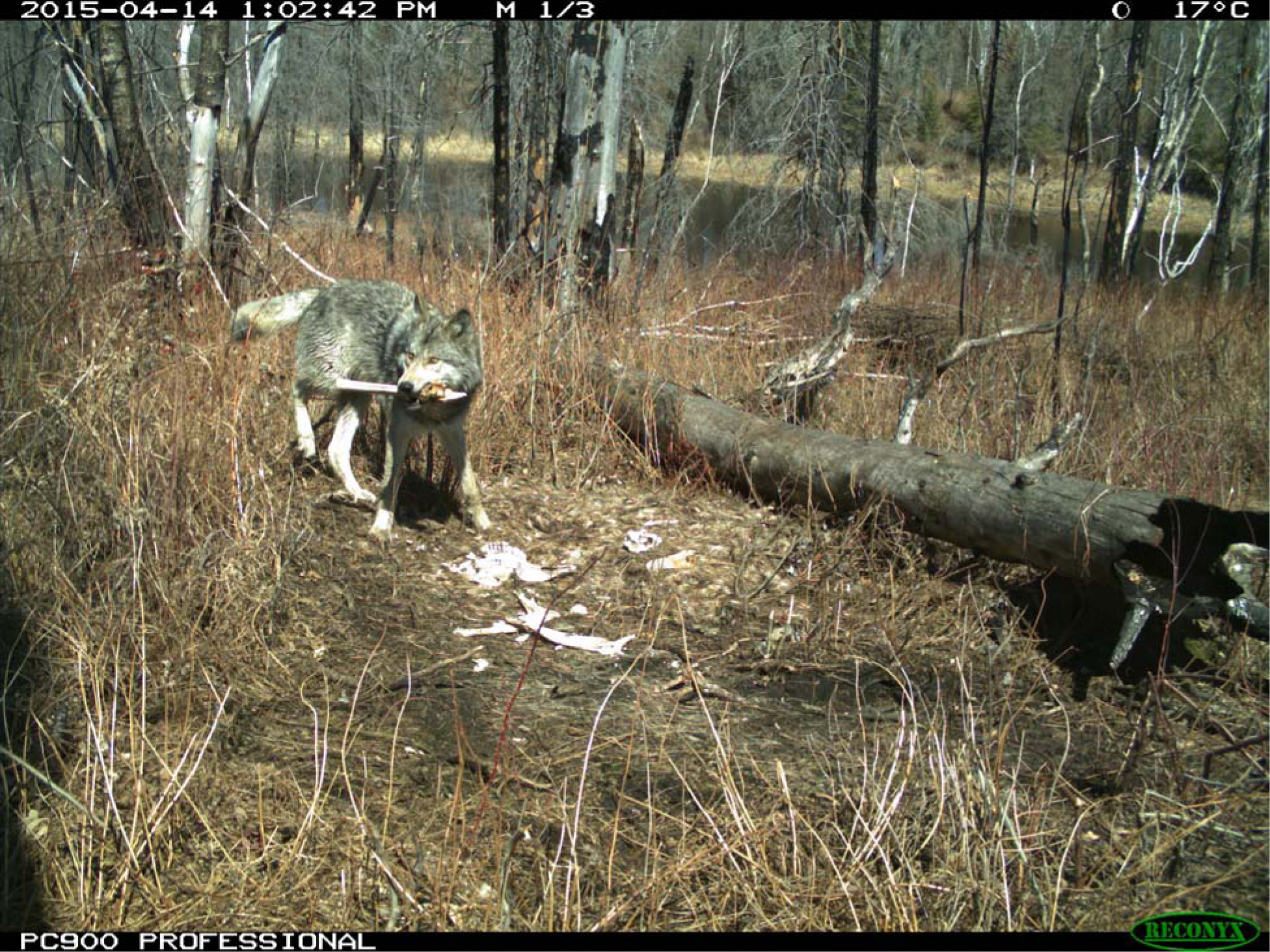
A wolf (*Canis lupus*) at a scavenging site of a hunter killed moose (*Alces alces*) visited by collared wolves. Reconyx camera deployed by Daniel Dupont north of Bissett during the first cluster investigation in October and picture captured in April 2015 when the wolves returned to the site again.

