## Supplementary material for "Evidence for optimal behavior of predators from parallel field investigations in two distinct wolf-prey systems": S1 Methods Details S2 Additional Results

### **Supplemental File**

### **S1. Methods Details**

Data and code (Python version 2.7.1, R version 4.3.0) and are available on GitHub <https://github.com/CMProkopenko/wolf_clusters> and will be archived on Zenodo.

**Glossary**

Cluster: Group of points, seed begins at 300m and 96 hours

Centroid: Centre of the cluster

Investigated Site: area visited during fieldwork to determine the behaviour occurring at clusters

Join: Combining the cluster data fields with the investigated data fields in a single file

Locations/Points: GPS locations from wolf collars

Merge: Horizontally combining the clusters or locations from individuals together

Match: when there is a spatial overlap in clusters with investigated sites

Sites: A spatial area corresponding to a behavior defined by physical evidence, can be composed of multiple clusters from one or more wolves.

**Data Cleaning Protocol**

Location downloads occurred frequently in GHA 26 (every second week) and RMNP (usually twice weekly). Sequential data downloads created segments of data that broke up large clusters and thus inflated the number of smaller clusters occurring in an area. The increase in small clusters did not change the outcomes of our observations site investigation but would change the characteristics of the cluster including radius, duration, first and last point, revisits etc. Following fieldwork, we downloaded all location data for each wolf and reran the cluster algorithm to get a comprehensive summary of clusters. We matched the investigated site locations recorded in the field to the new clusters with an overlap tolerance of 25 m and flagged data when investigations occurred early or late in reference to cluster formation time. We then reviewed site and cluster matches to determine the primary and secondary behavior and primary and secondary prey. These decisions were guided by which behavior was the primary motivation for occupying the area for a given time (Table 1 Main Text). A hierarchy was outlined to prioritize behaviors occurring at sites. For example, if a beaver kill overlapped with a den or rendez-vous, the overlap clusters were assigned the primary behavior of ‘den’ and a secondary behavior of ‘kill’. If overlapping clusters indicated a double kill (i.e. two prey individuals killed), then either the largest prey or the first location was assigned as the primary prey. The matched and joined file of all the sites was processed to identify the number of clusters and wolves at each site, first and last point of overlapping clusters, and total number of each prey species killed. A detailed workflow of steps is in the supplementary information and all code is available on GitHub.

**Niche Breadth**

We calculated wolf food niche breadth using Levins’ (1968) formula:

$B = \frac{1}{\Sigma{pi}^{2}}$,

where *p_i_* is the proportion of each food item that contributed to the total diet of wolves from both cluster and scat data. Prey species were classified as (1) adult ungulates; (2) neonate and calf ungulates; (3) medium-sized mammals (beavers, hares); and (4) others (bears, small rodents, plants, waste, etc.), which was composed of items that contributed less than 5% individually and 10% total to the overall wolf diet.

**Home Range Estimation**

Prior to calculating aKDEs, we visually assessed and removed any extra-territorial forays that deviated outside the general pattern of territorial space use. As well, we removed individual seasons that had fewer than 50 GPS locations (Dickie 2022). We then calculated pack-level seasonal home ranges by averaging individual seasonal home ranges, weighted by the density of points.

### **S2: Additional Results**

**Clusters**

Den sites were the largest clusters and had the longest residency time with a median of 36 locations and 166 hours in RMNP and 27 locations and 182 hours in GHA 26. Rendez-vous sites had fewer locations (median: RMNP = 25, GHA 26 = 23) and shorter median duration than den sites (RMNP = 152 hours, GHA 26 = 167 hours), however rendez-vous sites were spatially the largest with a median radius of 279 m in RMNP and 276 m in GHA 26. In comparison, resting clusters had some of the fewest locations, with a median of 7 locations for both study areas, and a duration of 14 hours for RMP and 16 hours for GHA 26. Wolves in GHA 26 had more clusters classified as revisits, but wolves spent a similar amount of time revisiting previous kills in both areas (Figure 2, Table S2.1).

**Table S2.1.** The median and range of cluster characteristics for each behavior by study area. Locations are the number of locations occurring at a cluster, if a cluster was left and returned to it would not count the locations outside of the cluster area. Duration of a cluster is the time from the first location of the cluster to the final location. Returns indicate when a wolf leaves the cluster for a few locations and then returns to have more locations included in the cluster. The locations away are not included in the ‘locations’ at cluster but are calculated in the cluster creation algorithm. The radius of the cluster area in meters that contains all locations in the cluster. The investigation delay is calculated as the investigation date subtracted from the final cluster date. Negative numbers indicate investigation happened after the cluster, while a positive number means the investigation occurred before or during the cluster. Early investigations can happen for multiple reasons; for example, long clusters with many revisits (e.g. dens, rendez-vous or resting sites), when a prey item was discovered before consumption or scavenging, or when one wolf from a pack visited before other wolves.

| **Behavior** | **Study Area (Total Count)** | **Locations** | **Duration (hours)** | **Returns (Count) and % Away** | **Radius (meters)** | **Investigation delay (days)** |
| --- | --- | --- | --- | --- | --- | --- |
| Den | RMNP (31) | 36 (3, 142) | 166 (4, 732) | 7 (0, 28)  55% (0, 82%) | 224 (30, 473) | -10 (-72, 4) |
|  | GHA 26 (105) | 27 (2, 260) | 182 (2, 1345) | 5 (0, 40)  30% (0, 88%) | 241 (15, 489) | -30 (-175, 37) |
| Rendez-vous | RMNP (60) | 25 (2, 135) | 152 (6, 785) | 4 (0, 26)  58% (0, 90%) | 279 (52, 506) | -15 (-55, 5) |
|  | GHA 26 (30) | 23 (3, 130) | 167 (4, 656) | 5 (0, 20)  58% (0, 92%) | 276 (69, 488) | -47 (-169, -3) |
| Moose Kill | RMNP (169) | 16 (2, 102) | 68 (2, 428) | 2 (0, 19)  48% (0, 94%) | 204 (5, 490) | -7 (-139, 4) |
|  | GHA 26 (164) | 22 (2, 178) | 92 (4, 674) | 3 (0, 17)  42% (0, 94%) | 247 (12, 479) | -28 (-173, 0) |
| Elk Kill | RMNP (67) | 11 (3, 41) | 60 (4, 204) | 2 (0, 8)  47% (0, 92%) | 240 (30, 430) | -10 (-46, 0) |
|  | GHA 26 (0) |  |  |  |  |  |
| Deer Kill | RMNP (64) | 8 (2, 26) | 22 (4, 122) | 1 (0, 5)  37% (0, 94%) | 195 (18, 471) | -6 (-51, 0) |
|  | GHA 26 | 13 (2, 66) | 47 (4, 450) | 2 (0, 17)  22% (0, 93%) | 210 (22, 523) | -19 (-148, 0) |
| Calf Kill | RMNP (21) | 9 (2, 50) | 26 (2, 142) | 1 (0, 5)  40%, 0, 85%) | 155 (24, 473) | -7 (-111, 3) |
|  | GHA 26 (92) | 16 (4, 72) | 63 (6, 366) | 1 (0, 7)  36% (0, 89%) | 233 (9, 439) | -16 (-146, 2) |
| Beaver Kill | RMNP (18) | 10 (5, 9) | 46 (10, 252) | 1 (0, 9)  50% (0, 90%) | 113 (10, 327) | -15 (-55, 3) |
|  | GHA 26 (175) | 7 (2, 61) | 22 (2, 280) | 0 (0, 11)  0% (0, 95%) | 141 (5, 510) | -20 (-270, 0) |
| Revisit | RMNP (48) | 6 (2, 80) | 19 (2, 280) | 0 (0, 7)  0% (0, 95%) | 188 (12, 397) | 0 (-42, 20) |
|  | GHA 26 (212) | 6 (2, 72) | 16 (2, 480) | 0 (0, 16)  0% (0, 96%) | 140 (1, 451) | -8 (-133, 96) |
| Probable Kill | RMNP (95) | 7 (2, 77) | 20 (2, 520) | 0 (0, 16)  0% (0, 93%) | 151 (6, 661) | -11 (-62, 0) |
|  | GHA 26 (23) | 6 (2,73) | 18 (2, 436) | 0 (0, 13)  15 (0, 95%) | 146 (3, 496) | -21 (-157, 4) |
| Scavenge | RMNP (89) | 8 (2, 57) | 38 (2, 980) | 1 (0, 20)  43% (0, 94%) | 184 (11, 506) | -7 (-250, 6) |
|  | GHA 26 (196) | 8 (2, 94) | 45 (2, 450) | 1 (0, 17)  54% (0, 95%) | 198 (5, 513) | -20 (-267, 52) |
| Resting | RMNP (226) | 7 (2, 69) | 14 (2, 428) | 0 (0, 12)  0% (0, 91%) | 121 (2, 402) | -9 (-63, 7) |
|  | GHA 26 (385) | 7 (2, 115) | 16 (2, 115) | 0 (0, 15)  0% (0, 95%) | 121 (1, 479) | -19 (-156, 6) |
| Other | RMNP (161) | 7 (2, 112) | 18 (2, 714) | 1 (0, 18)  31% (0, 94%) | 155 (1, 497) | -12 (125, 7) |
|  | GHA 26 (260) | 5 (2, 67) | 32 (2, 450) | 1 (0, 17)  46% (0, 96%) | 146 (3, 474) | -21 (-156, 7) |

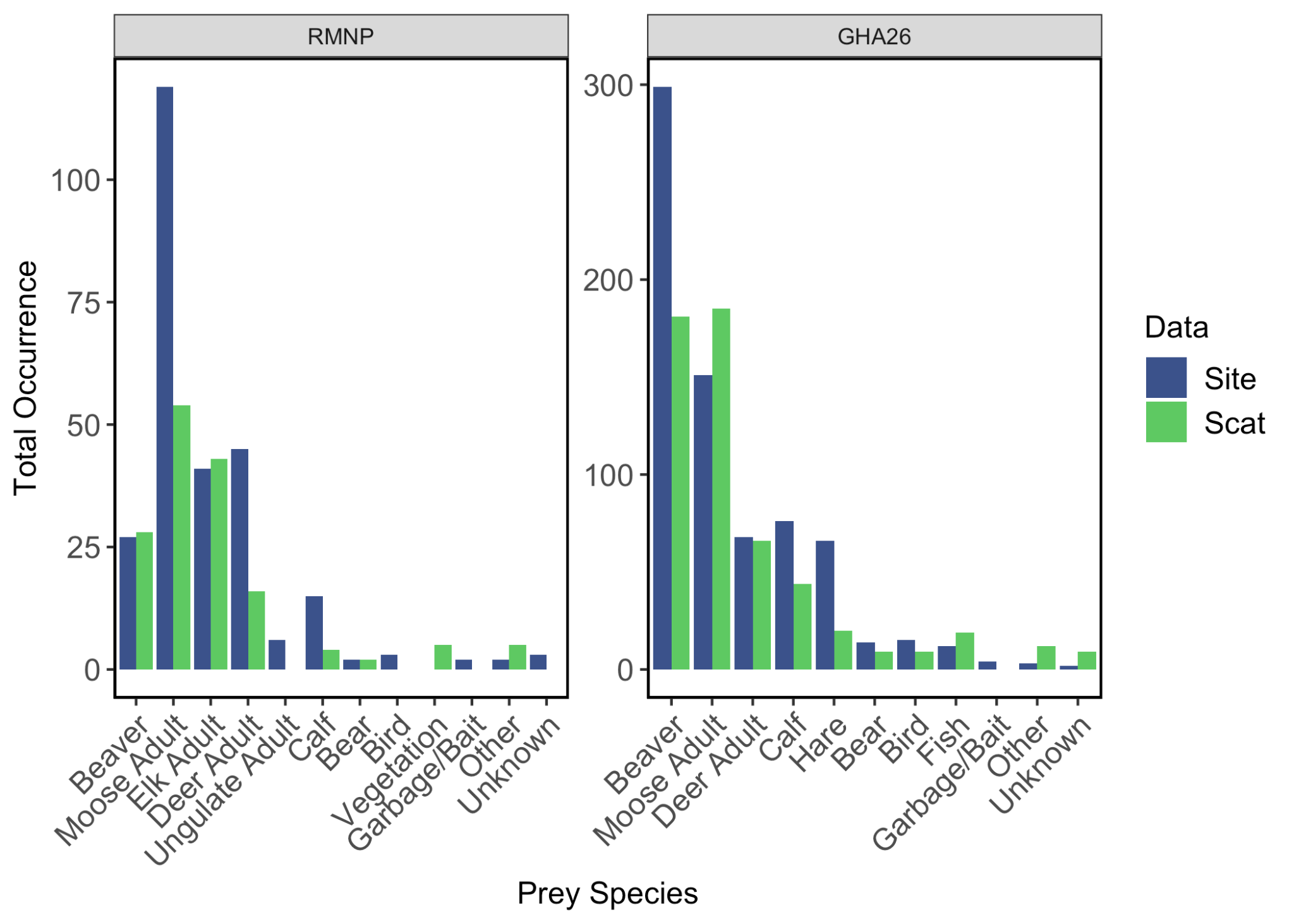

**Figure S2.1.** Occurrence of food source in gray wolf diets in RMNP (2016 - 2017) and GHA 26 (2014-2019) determined through two data sources; kill site investigations of clusters from collared wolves (i.e., Site) or found in wolf scat samples collected at kill sites (i.e., Scat). Data was collected year round; site results summarize diet composition found at kill, scavenging, and probable kill sites, while scat data was collected at all site types.

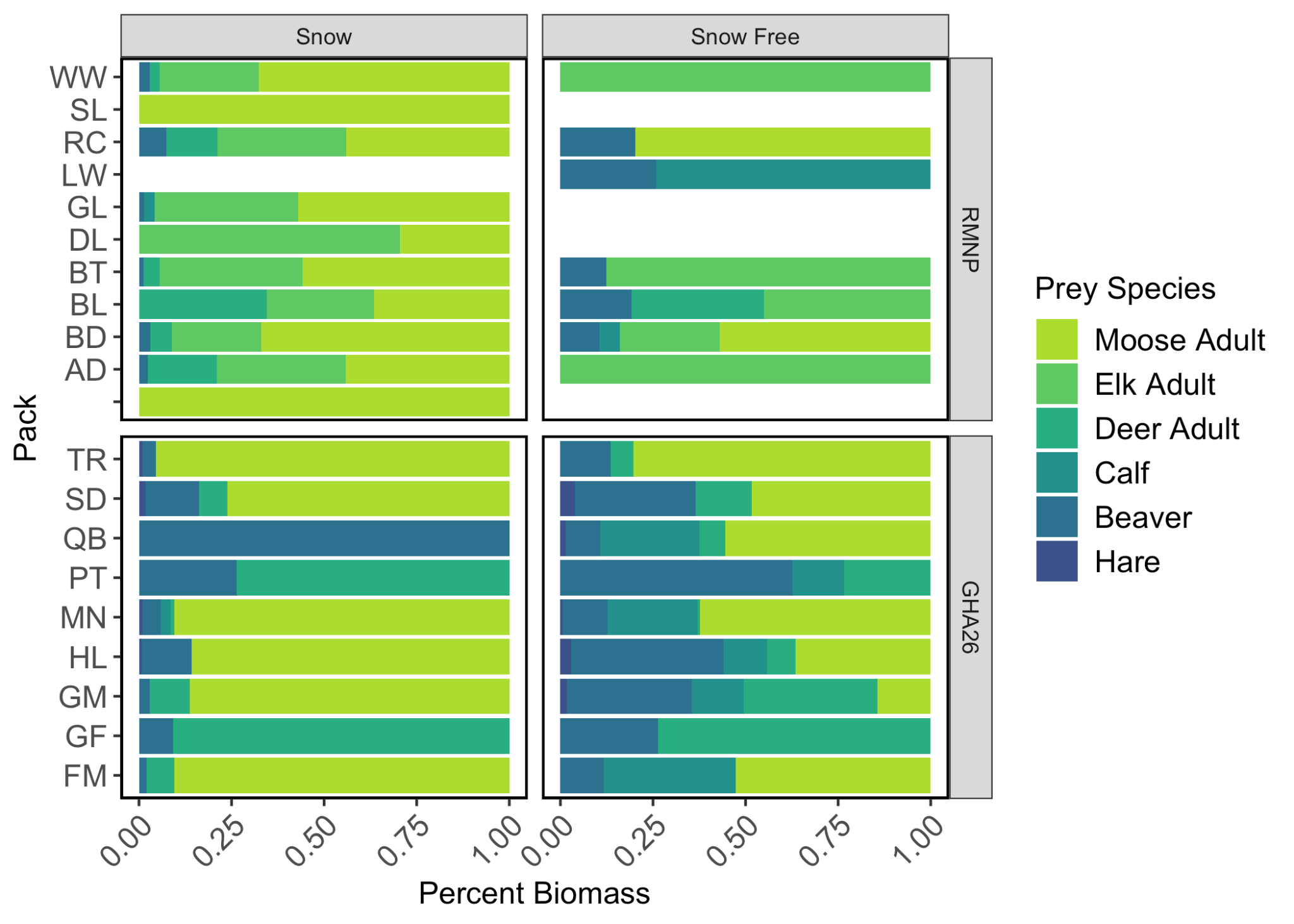

**Figure S2.2** Seasonal percent biomass contribution of prey type to gray wolf diet across packs in RMNP (2016 - 2017; n = 8) and GHA 26 (2014-2019; n = 13) determined through scat samples from collared wolves.

**Home Range Estimation**

From the 95% aKDE isopleths, we identified one wolf pack in RMNP that shifted their home range during the study period. We also identified two individual wolves in GHA 26 that left their original pack and spent the remainder of the study period traveling around the region and did not develop a stable home range. As a result, we removed the one wolf pack in RMNP and two individual wolves from GHA 26 and only reported home range sizes for wolf packs with stable home ranges.

**Table S2.2.** Seasonal (snow and snow free) and full year 95% aKDE home range sizes of wolf packs in Riding Mountain National Park and Game Hunting Area 26.

|  |  | Season | | |
| --- | --- | --- | --- | --- |
| **RMNP** | | Snow | Snow Free | Full Year |
| *2016* | |  |  |  |
|  | Baldy Lake | 173.92 | 121.48 | 137.94 |
|  | Deep Lake | 4116.69 | 1736.02 | 2621.67 |
|  | Gunn Lake | 300.33 | 192.94 | 300.12 |
|  | Whitewater | 289.25 | 285.17 | 460.49 |
| *2017* | |  |  |  |
|  | Audy | 18073.47 | 9269.74 | 14024.07 |
|  | Baldy Lake | 159.10 | 128.06 | 158.24 |
|  | Birdtail Valley | 358.48 | 219.30 | 303.36 |
|  | Ranch Creek | 194.48 | 67.41 | 139.12 |
|  | Spruce Lake | 312.32 | 212.81 | 301.90 |
| **GHA 26** | |  |  |  |
| *2014* | |  |  |  |
|  | Gem-Flintstone | 1626.38 | 3481.19 | 12901.34 |
|  | Quesnel-Bissett | 1260.73 | 1083.66 | 1377.29 |
| *2015* | |  |  |  |
|  | Frenchman | 1094.11 | NA | 1094.11 |
|  | Manigotagan | 1758.89 | 983.24 | 1324.02 |
| *2016* | |  |  |  |
|  | Frenchman | 1704.71 | 526.62 | 1725.93 |
|  | Happy Lake | 1445.46 | 865.60 | 1000.25 |
|  | Manigotagan | 934.45 | 616.10 | 727.38 |
|  | Pointe | 339.85 | 179.75 | 337.51 |
|  | Sandy River | 881.16 | 214.82 | 706.21 |
| *2017* | |  |  |  |
|  | Manigotagan | 3770.02 | 3478.97 | 2740.76 |
|  | Maskwa | 233.04 | 204.92 | 213.89 |
|  | Sandy River | 670.49 | 324.38 | 555.85 |
| *2018* | |  |  |  |
|  | Great Falls | 628.50 | 88.59 | 349.92 |
|  | Great Falls (lone wolf) | 8377.63 | 8881.95 | 9164.21 |
|  | Happy Lake | 478.18 | 231.30 | 458.07 |
|  | Manigotagan | 1067.50 | 353.73 | 972.98 |
|  | Sandy River | 685.19 | 606.32 | 705.52 |
|  | Snowshoe | 1939.78 | NA | 1939.78 |
|  | Turtle | 1243.46 | 4445.85 | 2568.68 |
| *2019* | |  |  |  |
|  | Black River (lone wolf) | 6627.33 | 21443.75 | 12241.28 |
|  | Frenchman | 398.44 | 281.35 | 382.31 |
|  | Happy Lake | 4076.43 | 2565.31 | 3758.43 |
|  | Sandy River | 448.16 | 240.25 | 319.09 |
|  | Smoky | 512.86 | 551.95 | 525.77 |
|  | Turtle | 924.77 | 876.36 | 853.09 |

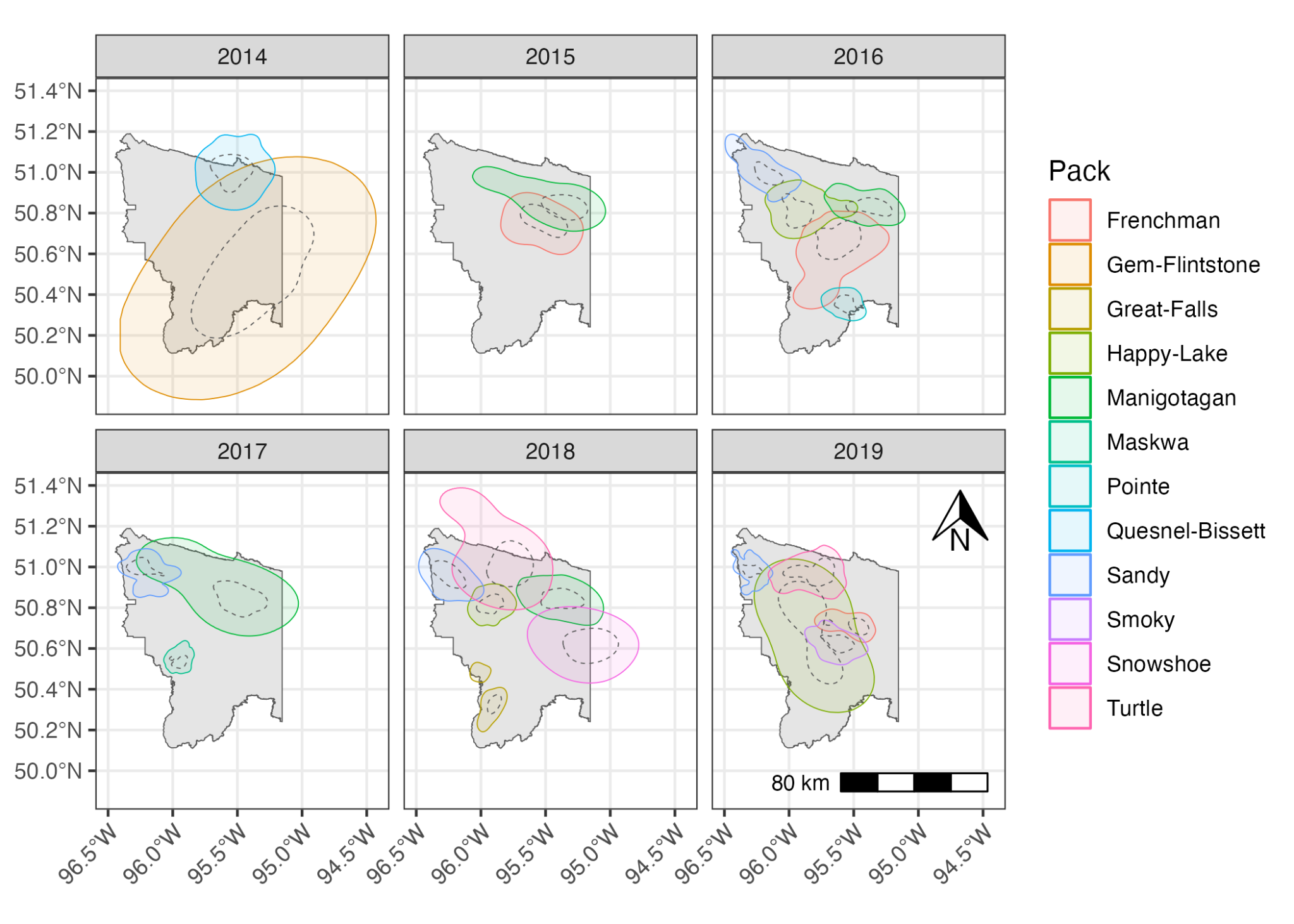

**Figure S2.3.** Full year 95% aKDE home range sizes of wolf packs (2014-2019; n = 13) in Game Hunting Area 26 by year. Solid lines are 95% isopleths and dashed lines are 50% isopleths of GPS collared wolves. Packs ranged from 1-3 collared individuals throughout the year; isopleths represent density weighted estimates of stable home range boundaries.

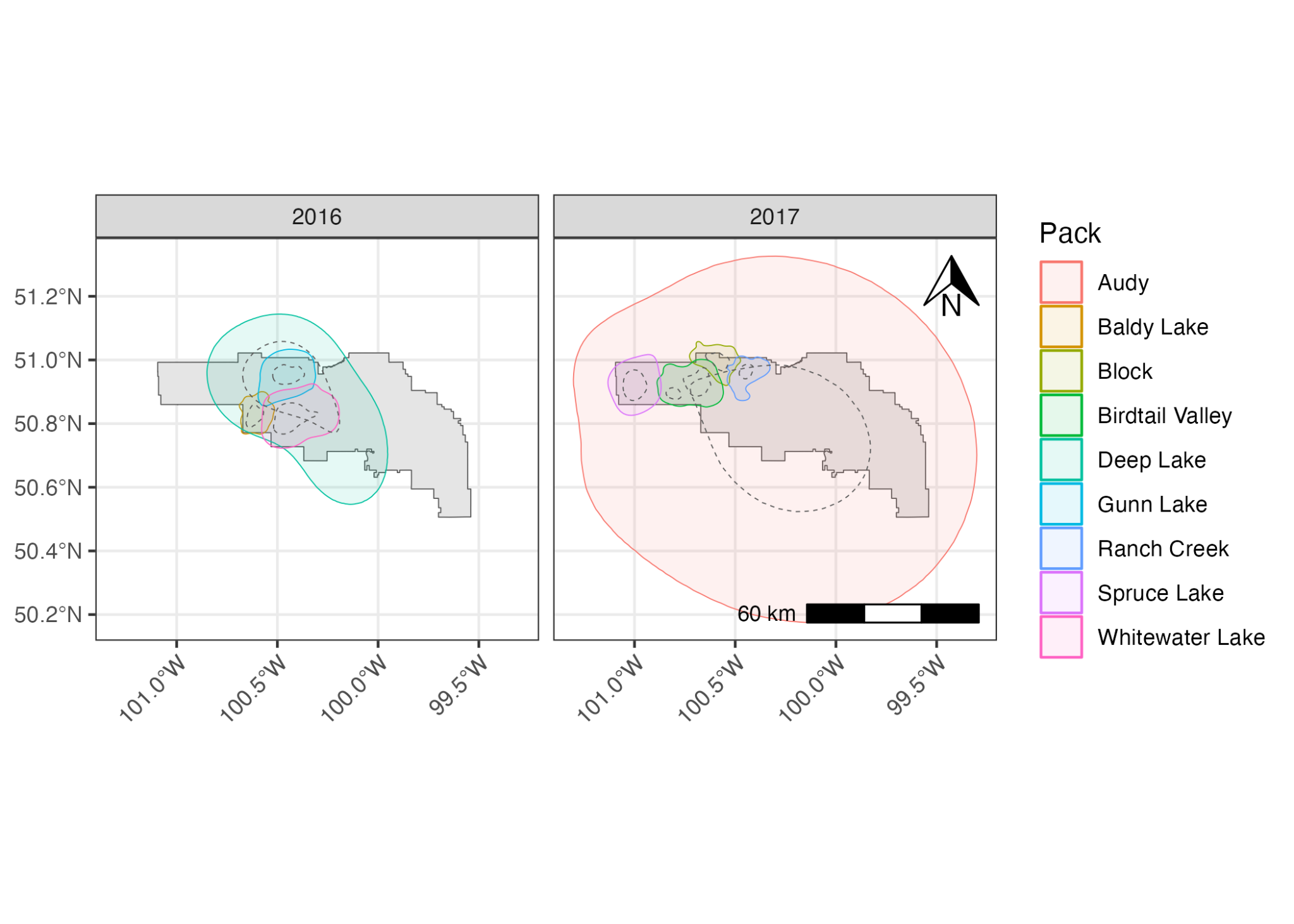

**Figure S2.4.** Full year 95% aKDE home range sizes of wolf packs (2017-2019; n = 9) in Riding Mountain National Park by year. Solid lines are 95% isopleths and dashed lines are 50% isopleths of GPS collared wolves. Packs ranged from 1-2 collared individuals throughout the year; isopleths represent density weighted estimates of stable home range boundaries.

**Step Length**

In general, GHA 26 wolves moved farther over the 2-hour time periods (median = 154 m, mean = 1077 m, range = 0 – 18028 m) than RMNP wolves (median = 125 m, mean = 793 m, range = 0 – 18710 m). GHA 26 wolves tended to move shorter distances in the snow season (median = 119 m, mean = 994 m, range = 0 – 18027 m) than the snow free season (median = 192 m, mean = 1152 m, range = 0 – 16023 m). RMNP wolves did not show as dramatic differences between the snow (median = 148 m, mean = 799 m, range = 0 – 14885 m) and snow free seasons (median = 108 m, mean = 788 m, range = 0 – 18710 m).

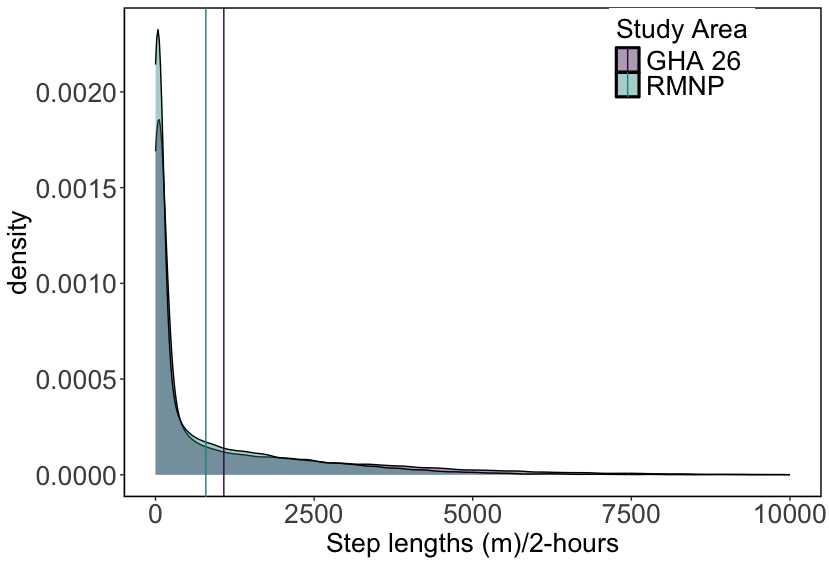

**Figure S2.5.** Step length distributions for wolves collared in GHA 26 (purple) and RMNP (teal green). Vertical lines indicate median step lengths. GHA 26 median = 154 m, mean = 1077 m, range = 0 – 18028 m; RMNP median = 125 m, mean = 793 m, range = 0 – 18710 m.

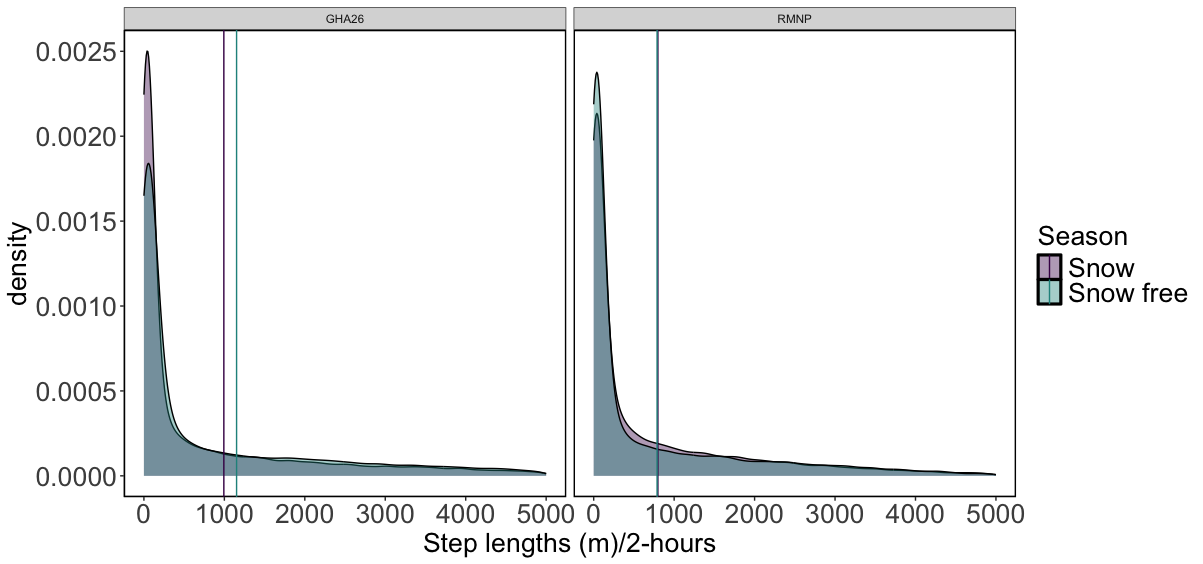

**Figure S2.6.** Step length distributions for GHA 26 and RMNP in the Snow (purple) and Snow free (teal) seasons. Vertical lines indicate median step lengths. GHA 26 in the snow season median = 119 m, mean = 994 m, range = 0 – 18027 m; snow free season (median = 192 m, mean = 1152 m, range = 0 – 16023 m). RMNP wolves snow season median = 148 m, mean = 799 m, range = 0 – 14885 m; and snow free seasons median = 108 m, mean = 788 m, range = 0 – 18710 m.
